# Effect of fumarate and live yeast on methane emissions, rumen fermentation, blood metabolites, and lactation performance in dairy goats

**DOI:** 10.1101/2024.08.31.610449

**Authors:** Huixin Dong, Jinlei Tan, Shuaishuai Li, Jingbo Ma, Guangfu Tang, Jiake Dai, Junhu Yao, Zongjun Li

## Abstract

The objective of this study was to evaluate the effects and combined effects of fumarate (32 g/d) and active dry yeast (ADY) (1.5 g/d) on methane emissions, rumen fermentation, microbial community and function, apparent total tract digestibility, blood metabolites, and lactation performance in 28 lactating goats using a randomized complete block design with a 2 × 2 factorial arrangement. Fumarate supplementation inhibited methane emission rates for 3 h post-feeding, while ADY increased the methane emission rates for 2 h pre-feeding, without combined effects between them. Fumarate increased the rumen pH and reduced total VFA concentration without changing the VFA profiles before feeding. Based on two custom dataset of 4,674 metagenome-assembled genomes and 7,758,615 nonredundant genes, neither fumarate nor live yeast influenced the rumen microbial community and function. The addition of fumarate reduced the concentrations of glucose, serum malondialdehyde (MDA), and ammonia in the serum, as well as the activity of insulin, while increasing the activity of serum malate dehydrogenase (MDH) and the levels of urea in both serum and milk. The addition of live yeast reduced serum glucose levels and increased serum MDA concentrations and MUN levels. There was a negative interaction between fumaric acid and live yeast on the levels of urea nitrogen in serum and milk.

**Conclusions:** The methane inhibition effect of fumaric acid persists for 3 h post-feeding, reflecting its retention or metabolism time in the rumen. Fumaric acid enhances the TCA cycle and the urea cycle in dairy goats, thereby improving energy utilization efficiency and reducing the risk of ammonia toxicity.

## Introduction

Methane emissions from ruminants not only contribute 9% to 11% of total anthropogenic greenhouse gas emissions, but also drain 2% to 12% of dietary energy (Johnson and Johnson, 1995; FAO, 2023). Recently, many additives, such as fumarate and 3-nitrooxypropanol, have well-documented methane mitigating effects (Li et al., 2018a; Maigaard et al., 2024). However, the accompanied effects of these additives, especially bioactive additives, on host metabolism also need be documented to be harmless to animal health, and preferably be beneficial.

Although the pathways of methanogenesis in the rumen has been fully understood, few strategies have proven effective in reducing methane emissions in a residue-free and non-toxic manner (Wang et al., 2023). Fumarate, as an intermediate in the succinate-propionate pathway of rumen microbes, has been widely exploited as an approach to redirect hydrogen away from methanogenesis to propionogenesis, which represents an energy-rendering pathway due to incorporating hydrogen energy and the main precursor of gluconeogenesis in ruminants (Li et al., 2018a; Maigaard et al., 2024). Up to now, little attention has been paid to the potential effects of fumarate supplementation on body metabolism of ruminants. Our previous findings in dairy goats showed that fumarate supplementation improved the blood total antioxidant capacity (T-AOC), but its underlying molecular mechanisms was unclear (Li et al., 2021c). Fumarate also is an intermediate in the citric acid cycle of animals (Wei et al., 2023), which intake could elevate the thermogenic respiration and T-AOC in mice. So, we hypothesized that part of the intaked fumarate by ruminant animals might escape from rumen fermentation, and be absorbed through the rumen wall and intestinal tract, and then influence the body metabolism.

Active dry yeast (ADY) of *Saccharomyces cerevisiae*, as a direct-fed probiotic microorganism for ruminants, is known to stabilize rumen environment and improve fiber digestion by de-oxygenation and stimulation of ruminal acetogenic, cellulolytic and lactate-utilizing bacteria (Fonty and Chaucheyras-Durand, 2006; Baker et al., 2022a). Both acetogenic and lactate-utilizing bacteria could compete hydrogen with methanogenesis, while cellulolytic bacteria supply hydrogen to methanogens, resulting opinions and resluts vary on the methane-suppressing effects of ADY (Lynch and Martin, 2002; Li et al., 2021a). Hence, the different stimulating extents of ADY on these bacteria need to compare. Additionally, fumarate and ADY, as a hydrogen oxidizer and an oxygen reducer, respectively, both could influence the ruminal redox environment, and thus likely have combined effect on the rumen fermentation and methanogenesis.

Therefore, the objective of this study was to systematically evaluate the effects and combined effects of fumarate and ADY on methane emissions, rumen fermentation, microbial community and function, apparent total tract digestibility, blood biochemical parameters, and lactation performance in lactating goats.

## MATERIALS AND METHODS

All experimental procedures were approved by the Northwest A&F University Animal Care and Use Committee (protocol number: DK2024065).

### Animals, diets, and experimental design

A total of 28 primiparous Guanzhong dairy goat with similar initial body weight (33 ± 2.7 kg) and days in milk (45 ± 13 d) were selected from same farm. The goats were grouped into seven blocks based on days in milk, body weight, and daily milk production (DMP). Animals within each block were randomly assigned to one of four dietary treatments: 1) control (no additive); 2) fumarate (Aladdin®, Shanghai, China) supplied at 34 g/d; 3) ADY (Alltech Inc., Nicholasville, KY) supplied at 1.5 g/d; 4) both fumarate and ADY supplied. The diets were provided as total mixed rations with a forage-to-concentration ratio of 48:52 (Table S1) and fed twice daily at 0730 and 1730 h. To ensure complete intake, the fumarate and/or ADY were top-dressed on a quarter of the provided rations and fed first. Goats were individually housed in 28 pens for three hours after feeding, during which the goats were milked. And Furthermore, goats were released to the barn after feed refusals was removed. Goats always had free access to water.

The goats have adapted to the environmental control chambers before feeding experiment. The feeding experiment lasted 38 d, including 28 d of adaptation to treatments and 10 d of sampling. Samples were collected as follows: measuring or collecting samples of dry matter intake (DMI), DMP, gas emissions, total digestive tract digestibility, and lactation performance on d 29-32; collecting blood samples on d 33-35; collecting rumen fluid samples on d 36-38.

### Measuring of methane emissions, apparent total tract digestibility and milk performance

The goats were moved from barn to chambers two days before gas measurements. Methane emissions of goats were measured using environmental chambers as previously described (Li et al., 2021c) during d 29-32.

To measure the DMI and apparent total tract digestibility of each goat, refusals and feces were collected and dried at 55°C for 72 h, and subsamples (about 100 g, wool removed) were ground through a 1 mm screen for further analysis. The dry matter (DM), crude protein (CP), neutral detergent fiber (NDF), and acid detergent fiber (ADF) contents of these samples were determined as previously described (Li et al., 2021c).

The DMP (morning and evening) of each dairy goat were recorded, and individual goat milk samples were collected. After being thoroughly mixed, 50 mL of the collected milk samples were stored in a refrigerator at 4℃ for subsequent analysis of milk components. Milk samples were analyzed for fat, protein, lactose, and milk urea nitrogen (MUN) within 24 h using an infrared milk analyzer (MilkoScan FT 120, FOSS, Hillerød, Denmark).

### Collection and analysis of blood samples and ruminal samples

Blood samples were collected from the external jugular vein into two of 10 mL blood collection tubes before morning feeding on d 33-35. The samples in the tubes were allowed to clot at room temperature for 30 min and then centrifuged at 3000 ×g for 15 min to obtain serum, which was stored at −80°C for subsequent analysis. The concentrations of serum malondialdehyde (MDA), glucose (GLU), ammonia, blood urea nitrogen (BUN), albumin (ALB), globulin (GLO), total protein (TP), beta-Hydroxybutyrate (BHBA), triglyceride (TG), low-density lipoprotein (LDL), high-density lipoprotein (HDL), malate dehydrogenase (MDH), pyruvate kinase (PK), alanine aminotransferase (ALT), aspartate aminotransferase (AST) were analyzed using respective commercial kits (Built Bioengineering Research Institute, Nanjing, China).

Rumen fluid samples were collected via oral tubing and manual vacuum pump before morning feeding during d 36-38 of the experiment. Approximately 50 mL of rumen fluid was discarded before sample collection to minimize saliva contamination. Rumen pH was immediately measured following sample collection. Rumen fluid were subsampled for the analysis of volatile fatty acids (VFAs) (5 mL with 1 mL 25% metaphosphoric acid added) and metagenomic sequencing (45 mL). The ruminal VFA concentrations were determined using gas chromatography (Agilent Technologies 7820A GC system, Palo Alto, CA, USA) as described by Li et al (Li et al., 2021c).

### Metagenomic sequencing and genome catalog constructing

The rumen fluid samples from each goat were freeze-dried and pooled. Microbial DNA was extracted following the protocol outlined by Yu and Morrison (Yu and Morrison, 2004), and deeply sequenced (62 Gb per sample on average) using the BGI DNBSEQ-T7 platform with 150-bp paired-end reads.

The metagenomic data was individually cleaned, assembled (Megahit), binned (metaBAT2, MaxBin2, and CONCOCT), and bin_refinement (-c 50 -x 10 options) using the metaWRAP pipeline (Uritskiy et al., 2018). The metagenomes of each block were co-assembled and co-binned as described above. The acquired metagenome-assembled genomes (MAGs) with CheckM (Parks et al. 2015) estimated genome quality score (completeness - 5*contamination + log(N50)) ≥50, completeness >50%, and contamination <10% were retained. To obtain a more complete and high-quality MAG catalog, we also collected 1,776 published MAGs from dairy goats in the same farm (Tan et al., 2024). These MAGs were dereplicated twice by <99% average nucleotide identity (ANI) to obtain the strain-level MAG catalog using dRep (Olm et al., 2017) with options --S_algorithm ANImf -sa 0.99 -nc 0.2 -comW 1 -conW 5 -N50W 1.

The MAGs were annotated for taxonomic classification using GTDB-Tk v2.0.0 (Chaumeil et al., 2020). The relative abundance of MAGs was quantified using the Quant_bins module of metawrap (Uritskiy et al., 2018). Relative abundance was expressed in CPM (copies per million reads), analogous to TPM (transcripts per million reads) in RNAseq analysis.

### Gene catalog construction and functional annotation

The Annotate_bins module of metawrap (Uritskiy et al., 2018) was utilized to cluster predicted genes using CD-HIT to construct a non-redundant gene catalog. Subsequently, functional annotation of metagenomes was conducted using eggNOG mapper (v2.1.7) in DIAMOND mapping mode DIAMOND software (V0.7.9) to blast unigenes to Kyoto Encyclopedia of Genes and Genomes (KEGG) database, Gene Ontology (GO) database, and Clusters of Orthologous Groups of proteins (COG) database (Huerta-Cepas et al., 2019). The relative abundance of genes in the samples was quantified using CoverM with parameters: --min-read-percent-identity 95, --min-read-aligned-percent 60, --min-covered-fraction 0.7, -m tpm.

### Statistical analysis

The average of repeated measurements (i.e., VFA and methane) for each goat was computed for statistical analysis. All data were analyzed as a repeated measures ANOVA using the PROC MIXED program in IBM SPSS Statistics 27. The statistical model included fumarate, ADY, and ADY × fumarate. Interactions as fixed effects, block groups as random errors. When there was ADY × fumarate interactions, differences among treatment means were assessed using the Tukey’s multiple comparison test.

Sample alpha diversity was estimated using Abundance-based Coverage Estimator (ACE), Shannon diversity index. Sample beta diversity was computed using Principal Coordinates Analysis (PCoA) based on Bray-Curtis dissimilarity (Bray and Curtis, 1957) in R v.4.3.2 (http://www.R-project.com). Multivariate analysis of variance was conducted using the PERMANOVA function from the R package vegan to compare the statistical differences in microbial composition between different experimental periods and treatments. Statistical significance was determined at *P* < 0.05, while tendency was declared at 0.05 ≤ *P* ≤ 0.10.

## Results

### Lactation performance, methane emissions, and apparent total tract digestibility

The effects of fumarate and ADY supplementation on DMI, lactation performance, methane emissions, and apparent total tract digestibility are shown in Table 1. There was no treatment effect on DMI, DMP, DMP/DMI, or apparent total tract digestibility. Both fumarate and ADY increased (*P* < 0.05) urea content in milk, and a negative interaction was found between them (*P* = 0.025).

**Table 1.** Effects of the dietary treatments on feed intake, milk performance, methane emissions, and dietary apparent nutrient digestibility of the dairy goats.

| Item | Treatment |  |  |  | SEM | P-value |  |  |
| --- | --- | --- | --- | --- | --- | --- | --- | --- |
|  | CON | FUM | ADY | FY |  | FUM | ADY | F×Y |
| DMI, kg | 1.62 | 1.65 | 1.64 | 1.69 | 0.054 | 0.452 | 0.595 | 0.847 |
| DMP, kg | 1.57 | 1.69 | 1.69 | 1.70 | 0.059 | 0.206 | 0.199 | 0.294 |
| DMP/DMI | 0.98 | 1.03 | 1.04 | 1.01 | 0.041 | 0.726 | 0.646 | 0.320 |
| Milk composition, % |  |  |  |  |  |  |  |  |
| Fat | 3.13 | 3.45 | 3.36 | 3.54 | 0.134 | 0.074 | 0.231 | 0.620 |
| Protein | 3.09 | 3.14 | 3.08 | 3.19 | 0.106 | 0.444 | 0.824 | 0.742 |
| TS | 11.8 | 12.0 | 11.8 | 12.2 | 0.244 | 0.186 | 0.635 | 0.637 |
| Lactose | 4.23 | 4.22 | 4.26 | 4.28 | 0.070 | 0.929 | 0.537 | 0.901 |
| Urea, mM | 0.34<sup>b</sup> | 0.43<sup>a</sup> | 0.45<sup>a</sup> | 0.42<sup>a</sup> | 0.022 | 0.187 | 0.039 | 0.025 |
| Methane emissions |  |  |  |  |  |  |  |  |
| CH<sub>4</sub>, L/d | 25.2 | 22.4 | 26.7 | 23.1 | 0.58 | 0.001 | 0.096 | 0.584 |
| CH<sub>4</sub>/DMI, L/kg | 16.7 | 12.1 | 16.7 | 14.1 | 0.72 | 0.007 | 0.915 | 0.994 |
| CH<sub>4</sub>/DMP, L/kg | 17.9 | 14.0 | 17.0 | 14.5 | 1.21 | 0.028 | 0.891 | 0.600 |
| Apparent nutrient digestibility, % |  |  |  |  |  |  |  |  |
| DM | 62.4 | 63.7 | 64.1 | 63.0 | 1.81 | 0.928 | 0.780 | 0.515 |
| NDF | 46.9 | 47.7 | 50.4 | 47.3 | 2.65 | 0.657 | 0.575 | 0.480 |
| ADF | 43.8 | 43.9 | 51.5 | 44.7 | 3.37 | 0.344 | 0.223 | 0.318 |
| CP | 81.0 | 81.4 | 81.8 | 81.0 | 1.03 | 0.850 | 0.855 | 0.520 |

<sup>a-b</sup>Means with different superscripts within a row differ ( $P < 0.05$ ).
CON, Control; FUM, Fumarate; ADY, Active dry yeast; FY, FUM+ADY; DMI, Dry matter intake; DMP, Daily milk production; TS, Total solids; DM, Dry matter; NDF, Neutral detergent fiber; ADF, Acid detergent fiber; CP, Crude protein; SEM, Standard error of means.

Fumarate supplementation suppressed (*P* < 0.05) methane emissions from goats, expressed as L/d (by 11.3%), L/kg DMI (by 27.4%), or L/kg DMP (by 21.8%). An interaction (*P* = 0.004) was detected between fumarate supplementation and post-feeding time for hourly methane emission rates (L/h), which was lower than control at the first 3 h post-feeding (Fig. 1). Goats fed ADY tended to have higher daily methane emissions (*P* = 0.096), with higher methane emission rates during the 11^th^ and 12^th^ h post-feeding. Hourly methane emission rates decreased with hours post-feeding (*P* < 0.001), but it increased in ADY-fed goats at the 10^th^ h after morning feeding and at the 8^th^ h after afternoon feeding compared with that of one hour before, likely attributing to intake the feed refusals. ADY supplementation had no effects on the methane yield (L/kg DMI or L/kg DMP).

**Figure 1.**
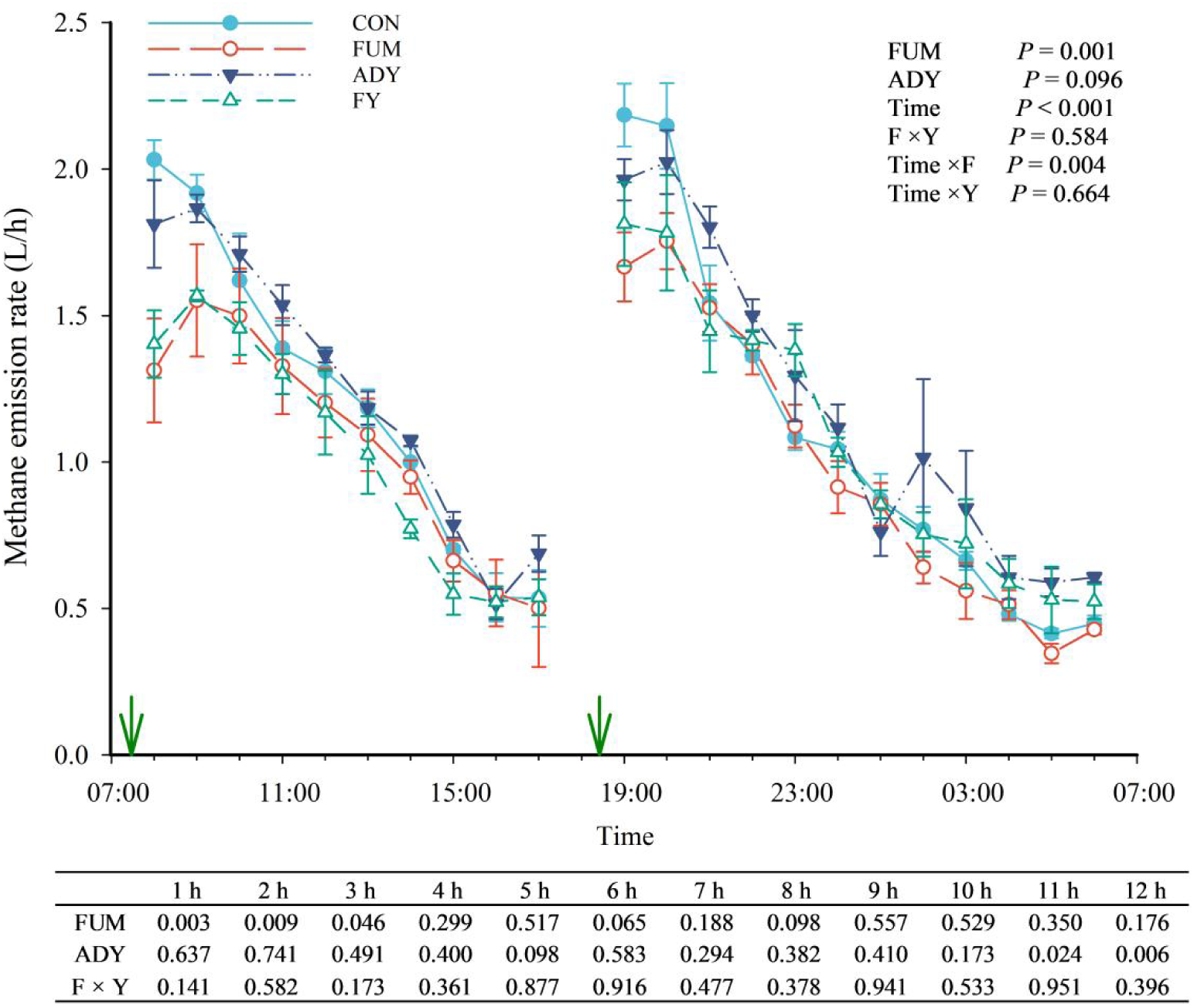
Effects of fumarate (FUM) and live active dry yeast (ADY) on diurnal methane emission rate (L/h, mean ± standard error) in dairy goats. CON, Control; FY, FUM+ADY.

### Rumen fermentation parameters, microbial community and function

Fumarate supplementation increased ruminal pH and reduced total VFA concentration without changing the VFA profiles (Table 2). Supplementation with ADY increased the molar proportion of butyrate (*P* = 0.039) without changing ruminal pH and VFA concentration.

**Table 2.** Effect of dietary treatments on rumen fermentation parameters.

| Item | Treatment |  |  |  | SEM | P-value |  |  |
| --- | --- | --- | --- | --- | --- | --- | --- | --- |
|  | CON | FUM | ADY | FY |  | FUM | ADY | F×Y |
| pH | 6.82 | 6.98 | 6.77 | 7.04 | 0.067 | 0.004 | 0.983 | 0.377 |
| VFA concentration, mM | 83.8 | 76.8 | 88.2 | 69.5 | 3.13 | <0.001 | 0.649 | 0.076 |
| Individual VFA molar proportion, % |  |  |  |  |  |  |  |  |
| Acetate | 65.4 | 65.5 | 65.6 | 64.9 | 0.57 | 0.632 | 0.730 | 0.555 |
| Propionate | 18.7 | 18.1 | 17.2 | 17.8 | 0.48 | 0.963 | 0.086 | 0.271 |
| Butyrate | 11.9 | 12.1 | 13.2 | 12.8 | 0.44 | 0.811 | 0.039 | 0.567 |
| Valerate | 1.32 | 1.31 | 1.30 | 1.36 | 0.803 | 0.882 | 0.732 | 0.976 |
| Isobutyrate | 1.19 | 1.36 | 1.24 | 1.45 | 0.106 | 0.086 | 0.499 | 0.861 |
| Isovalerate | 1.48 | 1.67 | 1.50 | 1.69 | 0.125 | 0.144 | 0.857 | 0.982 |
| A:P | 3.51 | 3.63 | 3.82 | 3.66 | 0.106 | 0.849 | 0.118 | 0.189 |

<sup>a-b</sup>Means with different superscripts within a row differ ( $P < 0.05$ ).
CON, Control; FUM, Fumarate; ADY, active dry yeast; FY, FUM+ADY; A:P, Acetate: propionate ratio; SEM,
Standard error of means.

After concatenation and quality filtering, a total of custom 4,674 MAGs, classified into 28 phyla, 39 classes, 80 orders, 137 families, and 513 genera. At the phylum level, predominant taxa included Bacteroidota (36.2%), Firmicutes_A (34.9%), Firmicutes_c (10.0%), and Firmicutes (7.3%) (Fig. 2D). Based on Bray–Curtis beta diversity, fumarate and ADY, either alone or in combination, had no impact on the microbial community and function (permutational multivariate ANOVA: *P* = 0.117, R^2^ = 0.121; function: *P* = 0.238, R^2^ = 0.117, Fig. 2A and Fig. 2B). Supplementation with fumarate and yeast had no effect on the proportion of carbohydrate enzyme genes ( Fig. 2C). The phylum Fibrobacterota was increased (P <0.05) by fumarate (Table 3). Within the genera, the relatibe abundances of the *Megasphaera* was increased (*P* < 0.05) by fumarate, and the genera *Lactobacillus* tended to gain higher relatibe abundance.

**Figure 2.**
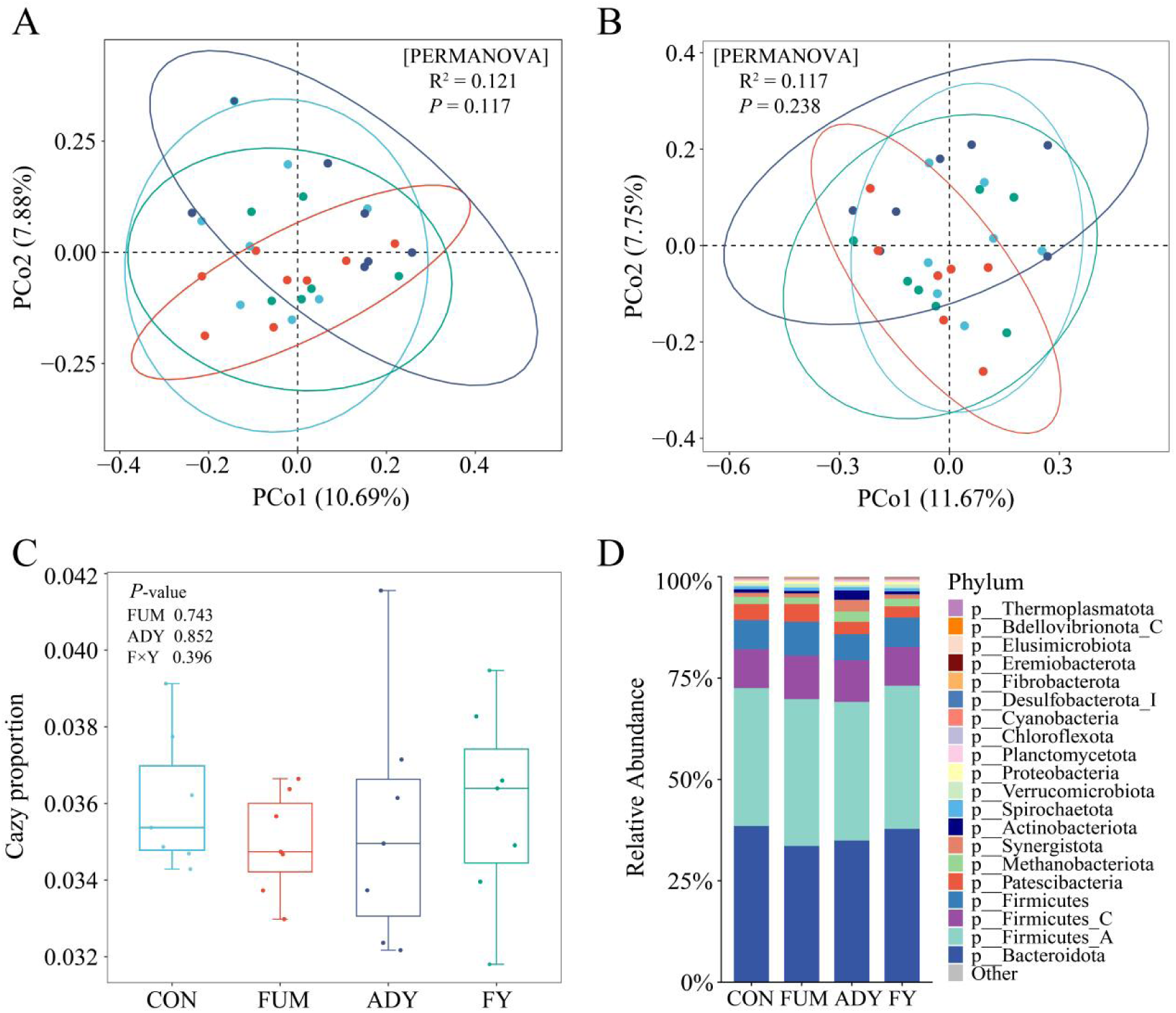
Principal Coordinate Analysis (PCoA) of rumen microbial community dissimilarities (A) and gene dissimilarities (B) between different treatments based on Bray-Curtis indices. Ellipses surrounding each treatment group represent a 95% confidence interval. (C) Box plot showing the proportion of carbohydrate enzyme genes relative to the total. (D) Stacked bar chart classifying species at the phylum level, with the top 20 abundance displayed. CON, Control; FUM, Fumarate; ADY, Active dry yeast; FY, FUM+ADY.

**Figure 3.**
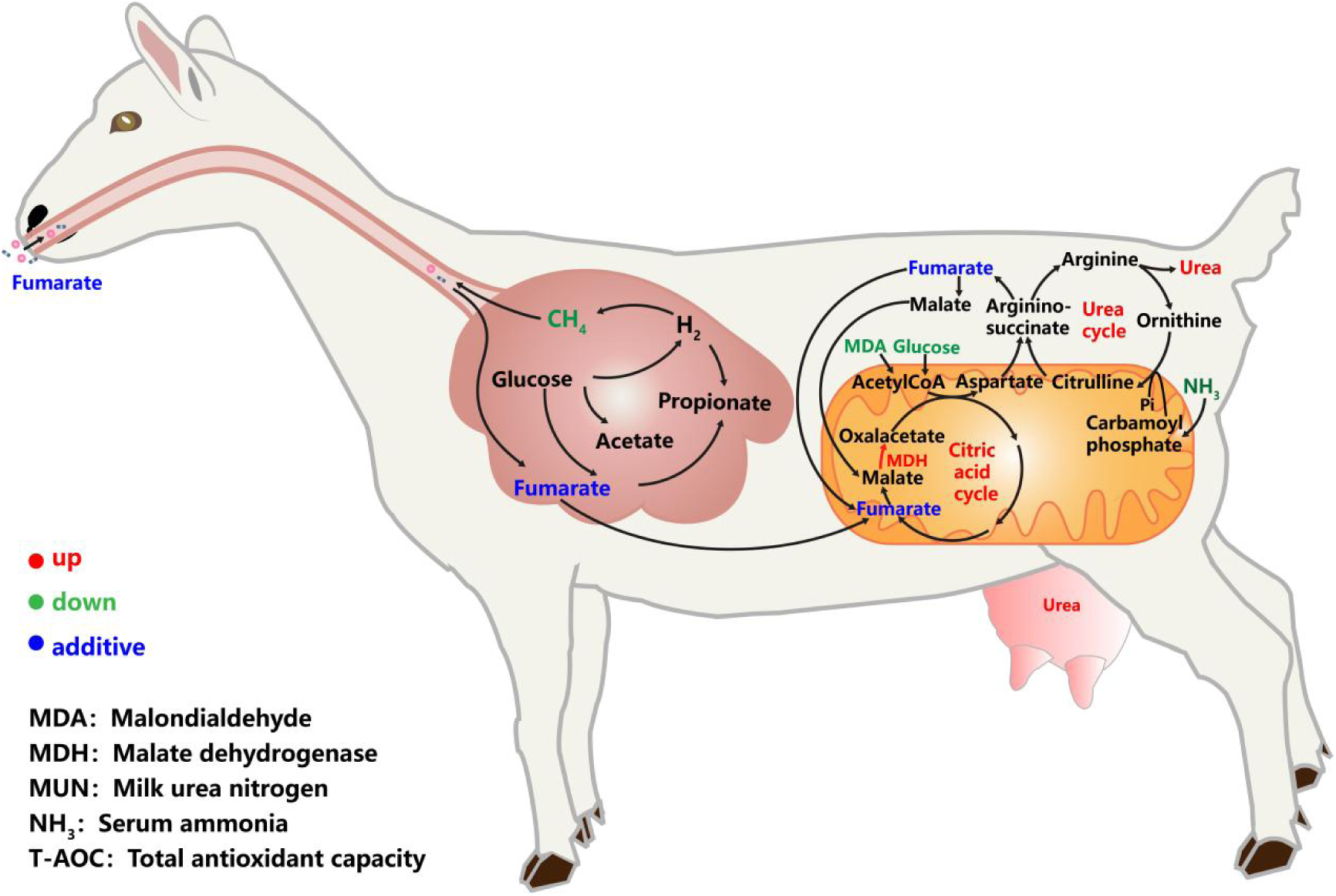
Mechanism of the action of Fumarate on the body of dairy goats.

**Table 3.** Effects of the dietary treatments on relative abundances of ruminal concerned bacterial and alpha diversity of microbial community.

| Item | Treatment |  |  |  | SEM | P-value |  |  |
| --- | --- | --- | --- | --- | --- | --- | --- | --- |
|  | CON | FUM | ADY | FY |  | FUM | ADY | F×Y |
| Ratio of *Saccharomyces cerevisiae* reads, ×10<sup>4</sup> | 0.93 | 0.64 | 0.38 | 0.87 | 0.106 | 0.849 | 0.118 | 0.189 |
| Relative abundances of concerned bacteria, CPM |  |  |  |  |  |  |  |  |
| *p__Fibrobacterota* | 11.6 | 18.8 | 8.08 | 13.3 | 2.915 | 0.044 | 0.140 | 0.742 |
| *p__Halobacteriota* | 0.09 | 0.15 | 0.16 | 0.16 | 0.058 | 0.632 | 0.502 | 0.603 |
| *p__Methanobacteriota* | 181 | 167 | 255 | 195 | 37.1 | 0.329 | 0.182 | 0.536 |
| *p__Thermoplasmatota* | 2.07 | 3.34 | 2.99 | 3.55 | 0.811 | 0.270 | 0.495 | 0.659 |
| *g__Megasphaera*, ×10<sup>-3</sup> | 7.62 | 15.2 | 3.43 | 16.3 | 3.80 | 0.011 | 0.681 | 0.482 |
| *g__Selenomonas_A* | 2.96 | 2.56 | 1.66 | 3.73 | 1.121 | 0.460 | 0.954 | 0.282 |
| *g__Selenomonas_B* | 26.7 | 25.1 | 12.7 | 34.6 | 8.71 | 0.255 | 0.794 | 0.191 |
| *g__Selenomonas_C* | 0.03 | 0.05 | 0.04 | 0.07 | 0.012 | 0.147 | 0.285 | 0.584 |
| *g__Acetobacter* | 0.07 | 0.05 | 0.34 | 0.06 | 0.178 | 0.215 | 0.234 | 0.282 |
| *g__Butyrivibrio* | 104 | 113 | 103 | 123 | 17.0 | 0.782 | 0.402 | 0.730 |
| *g__Lactobacillus* | 0.27 | 0.31 | 0.18 | 0.19 | 0.047 | 0.052 | 0.622 | 0.753 |
| *g__Prevotella*, ×10<sup>-3</sup> | 1.85 | 1.21 | 1.91 | 1.71 | 0.358 | 0.255 | 0.646 | 0.793 |
| Alpha diversity of microbial community |  |  |  |  |  |  |  |  |
| ACE, ×10<sup>4</sup> | 1.71 | 1.67 | 2.04 | 1.63 | 0.096 | 0.027 | 0.158 | 0.064 |
| Shannon | 7.00 | 7.06 | 6.67 | 7.09 | 0.127 | 0.070 | 0.246 | 0.153 |

CON, Control; FUM, Fumarate; ADY, Active dry yeast; FY, FUM+ADY; SEM, Standard error of means.

### Blood biochemical parameters

Supplementation of fumarate reduced (*P* < 0.05) the concentrations of ammonia, glucose and the activity of insulin (*P* = 0.025), and increased the the activity of MDH (*P* = 0.006) in blood (Table 4). Interactions (*P* < 0.05) between fumarate and ADY were observed for MDA, and urea in blood. Both fumarate and ADY reduced (*P* < 0.05) the concentrations of MDA and urea in blood. Supplementation of ADY reduced the concentrations of glucose.

**Table 4.** Effect of dietary treatments on blood metabolites of dairy goats.

| Item | Treatment |  |  |  | SEM | P-value |  |  |
| --- | --- | --- | --- | --- | --- | --- | --- | --- |
|  | CON | FUM | ADY | FY |  | FUM | ADY | F×Y |
| Ammonia metabolism |  |  |  |  |  |  |  |  |
| ALT, U/L | 9.67 | 8.86 | 6.25 | 9.01 | 1.320 | 0.470 | 0.230 | 0.191 |
| AST, U/L | 21.7 | 26.5 | 25.3 | 25.9 | 2.53 | 0.299 | 0.569 | 0.423 |
| Ammonia, μmol/L | 63.1 | 61.6 | 67.6 | 55.6 | 3.21 | 0.045 | 0.815 | 0.116 |
| Urea, mM | 5.80<sup>b</sup> | 6.70<sup>a</sup> | 6.69<sup>a</sup> | 6.40<sup>a</sup> | 0.162 | 0.077 | 0.085 | 0.001 |
| ALB, g/L | 27.9 | 27.0 | 26.0 | 26.9 | 0.62 | 0.995 | 0.129 | 0.158 |
| GLO, g/L | 39.8 | 41.5 | 42.8 | 40.7 | 1.95 | 0.908 | 0.578 | 0.338 |
| TP, g/L | 67.7 | 68.4 | 68.8 | 67.6 | 1.88 | 0.906 | 0.944 | 0.595 |
| A:G | 0.71 | 0.67 | 0.62 | 0.67 | 0.039 | 0.899 | 0.230 | 0.258 |
| Carbohydrate metabolism |  |  |  |  |  |  |  |  |
| MDH, U/mL | 0.55 | 0.66 | 0.59 | 0.68 | 0.033 | 0.006 | 0.336 | 0.841 |
| PK, U/L | 92.1 | 163 | 89.1 | 148 | 50.69 | 0.211 | 0.859 | 0.907 |
| Insulin, mIU/L | 29.2 | 18.7 | 24.2 | 20.3 | 3.02 | 0.025 | 0.578 | 0.279 |
| Glucose, mM | 3.38 | 3.11 | 3.15 | 3.03 | 0.071 | 0.010 | 0.037 | 0.291 |
| Lipid metabolism |  |  |  |  |  |  |  |  |
| T-chol, mM | 2.23 | 2.22 | 2.19 | 2.20 | 0.164 | 1.000 | 0.863 | 0.928 |
| MDA, μmol/L | 1.17<sup>a</sup> | 0.60<sup>b</sup> | 0.51<sup>b</sup> | 0.55<sup>b</sup> | 0.128 | 0.049 | 0.010 | 0.027 |
| BHBA, μmol/L | 205 | 198 | 226 | 200 | 13.0 | 0.215 | 0.368 | 0.474 |
| TG, mM | 0.24 | 0.21 | 0.22 | 0.22 | 0.022 | 0.598 | 0.689 | 0.555 |
| LDL, mM | 0.52 | 0.55 | 0.51 | 0.49 | 0.058 | 0.985 | 0.557 | 0.658 |
| HDL, mM | 1.41 | 1.43 | 1.35 | 1.29 | 0.166 | 0.890 | 0.555 | 0.804 |

<sup>a-b</sup>Means with different superscripts within a row differ ( $P < 0.05$ ).
CON, Control; FUM, Fumarate; ADY, active dry yeast; FY, FUM+ADY; T-AOC, Total antioxidant capacity; SOD, Superoxide dismutase; GSH-Px, Glutathione peroxidase; MDA, Malondialdehyde; ALT, Alanine aminotransferase; AST, Aspartate aminotransferase; BUN, Blood urea nitrogen; ALB, Albumin; GLO, Globulin; TP, Total protein; A:G, ALB: GLO ratio; MDH, Malate dehydrogenase; PK, Pyruvate kinase; GLU, Glucose; T-chol, Total cholesterol; BHBA, Beta-Hydroxybutyrate; TG, Triglyceride; LDL, Low-density lipoprotein; HDL, High-density lipoprotein; SEM, Standard error of means.

## Discussion

### Effects of fumarate supplementation on methane emissions

Theoretically, converting all 34 g of fumarate (0.29 mol) to propionate could potentially accept 0.29 mol hydrogen and reduce daily methane emissions by 1.8 L (Li et al., 2018b), which was lower than the actual decrease in the current study (average 2.8 L). Similar results have also been observed in our previous results (Li et al., 2018b, 2021c), suggesting that the mechanism of fumarate action in inhibiting methanogenesis is not solely an electron acceptor competing for hydrogen (Asanuma and Hino, 2000; Iwamoto et al., 1999). As fumarate supplementation did not affect DMI, the apparent nutrient digestibility, rumen microbial community composition or function of the dairy goats, indicating that fumarate was not by inhibiting feed intake, feed degradation or altering microbial community structure to reduce methane emissions. Therefore, we hypothesize that fumarate, as an intermediate in the propionate pathway, not only competes with methane for hydrogen but also stimulates the transcription or metabolic activity of the pathway, enhancing propionate fermentation from other substrates.

Yet, the decrease in methane emission rate (L/h) upon fumarate supplementation lasted for about 3 hours after feeding, consistent with a previous study (Maigaard et al., 2024), reflecting the ruminal retention or metabolism time of intaked fumarate that the intaked fumarate either washed out from rumen or converted to succinate within 3 hours from dosing. However, this study also found no significant difference in the proportion of propionate before feeding or 3 hours after feeding, which could explain why the propionate concentration in our results did not increase, which may be related to the timing of rumen fluid collection (before morning feeding). Additionally, according to the results of microbial community analysis, the addition of fumarate and yeast only had a short-term impact on the microbial community but did not affect microbial colonization, thereby potentially influencing rumen microbial fermentation processes. In summary, whether fumarate further inhibits methane emissions by enhancing the propionate fermentation pathway is a hypothesis that still requires further verification.

### Effects of fumarate on ruminal pH

*Fibrobacter succinogenes*, *Selenomonas ruminantium (ruminantium)*, and *Selenomonas ruminantium (lactilytica*) are fumarate-utilizing bacteria that compete with methanogens for H_2_ utilization (Castillo et al., 2004). which may be one of the reasons for the observed increase in ruminal pH.

Previous studies have shown that the addition of fumarate can lead to an increased lactate uptake, thereby reducing ruminal lactate accumulation and increasing pH (Araújo et al., 2011). *Megasphaera elsdenii* and *Selenomonas ruminantium* are also major lactate-utilizing species in the rumen (Henning et al., 2010). Among them, *Megasphaera elsdenii* is more significant because it can utilize 0.65-0.95% of the lactate in the rumen, thus competing with lactate producers for substrates (Counotte and Prins, 1981). Furthermore, *Megasphaera elsdenii* as a commensal group within the rumen of ruminants, are commonly added to feed as a direct-fed microbial (DFM) to help mitigate ruminal acidosis (Long et al., 2014; Weimer et al., 2015), which may be the second reason for the increase in pH.

VFAs are highly lipophilic compounds in their undissociated states and can easily traverse cell membranes (Aschenbach et al., 2011). When the intracellular pH is more alkaline than the extracellular pH. They can cross the cell membrane, dissociate, and accumulate within the cell (Russell et al., 1992). VFAs like fumarate may also cross the rumen wall and reach the hindgut, where they may further contribute to an increase in pH. This might also explain why pH is negatively correlated with the total concentration of volatile fatty acids (EL Packer et al., 2011), which is consistent with previous research findings (Li et al., 2018a; De Nardi et al., 2014).

### Effects of ADY supplementation on methane emissions

The impact of ADY on methane production varies across studies. (Mwenya et al., 2004) reported that adding ADY reduced methane emissions. (Bayat et al., 2015) observed no effect of ADY on methane emissions, and (Li et al., 2021b) found no impact on methane emissions when supplementing different doses of ADY. In summary, in this experiment, methane production, when expressed in grams per day, showed no differences, aligning with previous studies that found no effect of ADY on methane production (Bayat et al., 2015; Li et al., 2021b). Therefore, the ability of ADY to reduce methane production appears to be influenced by various factors.

Both acetogenic and lactate-utilizing bacteria may compete with methanogens for hydrogen, while cellulolytic bacteria supply hydrogen to methanogens (Baker et al., 2022b). As a result, opinions and findings on the methane-suppressing effects of ADY vary. In this experiment, ADY did not affect the microbial community, and consequently, had no impact on methane production or ruminal pH.

### Effects of fumarate on blood parameters

Assuming that fumarate quickly associates with the liquid fraction in the rumen, where the passage rate out of the rumen is fast (15%/h; (Krämer et al., 2013)), this rapid passage rate suggests that fumarate can be swiftly absorbed by the body and transported via the bloodstream throughout the system. Consequently, despite its short retention time in the rumen, fumarate can effectively enter the bloodstream and exert its antioxidant and metabolic regulatory functions within the body.

Fumarate serves as a crucial intermediate in both the TCA cycle and the urea cycle (Wei et al., 2023). MDA is the principal and most studied product of polyunsaturated fatty acid peroxidation, addition of fumarate salts can enhance the body’s antioxidant defense system, consistent with previous research findings (Del Rio et al., 2005; Meng et al., 2023). Glucose is a crucial substrate in the tricarboxylic acid cycle, providing energy to the animal body through glycolysis (Fernie et al., 2004). Several research suggests that there is a positive correlation between changes in blood glucose and insulin concentrations, and dietary supplementation response can alter the glucose-insulin system (Muñoz-Gutiérrez et al., 2005; Scaramuzzi et al., 2006). MDH widely distributed across various organisms, primarily participates in the TCA and catalyzes the reversible conversion between malate and oxaloacetate (Ge et al., 2010). In this experiment, the addition of fumarate may have promoted the TCA cycle, thereby reducing the levels of glucose and MDA in the blood and increasing MDH activity.

Proteins and non-protein nitrogen in feed are degraded in the rumen to produce ammonia. A portion of this ammonia is absorbed by the rumen wall, and after passing through the liver via the urea cycle, it eventually forms urea (Chikhou et al., 1993). Nitrogen stored in the form of urea is far less toxic than ammonia at equivalent concentrations (Wright et al., 1995). Consequently, the level of BUN serves as an indicator of protein deposition and is also an important parameter reflecting the functionality of both the kidneys and the liver (Stewart, 1977). In this study, supplementing with fumarate may have promoted the urea cycle, thereby reducing ammonia concentrations in the blood and increasing the production of BUN..The concentration of MUN is strongly linearly correlated with BUN (Hof et al., 1997), a point that can also be evidenced in the elevation of MUN.

In summary, fumarate, as an intermediate in both the tricarboxylic acid cycle and the urea cycle, may enhance the activity of these cycles, thereby reducing the risk of negative energy balance and ammonia toxicity, and exerting no adverse effects on the metabolism of dairy goats.

## Conclusions

Supplementing with fumarate is a promising approach to reducing methane emissions, with its inhibitory effect on methane persisting for 3 hours post-feeding. Fumarate enhances the TCA cycle and the urea cycle in dairy goats, thereby improving energy utilization efficiency and reducing the risk of ammonia toxicity.

## Acknowledgments

This work was supported by National Key Research and Development Program of China (award number: 2023YFE0111800), and the National Natural Science Foundation of China (award number: 31902126). We thank the High-Performance Computing Platform of Northwest A&F University and Computing Center in Xi’an for providing computing resources.

---

**Table S1.** Ingredients and chemical composition of the experimental diet.

| Item | Dry matter % |
| --- | --- |
| Ingredients |  |
| Corn silage | 21.2 |
| Alfalfa hay | 30.7 |
| Ground corn | 22.8 |
| Soybean meal | 6.5 |
| Corn germ meal | 3.1 |
| Cottonseed meal | 4.9 |
| Wheat bran | 8.1 |
| NaHCO<sub>3</sub> | 0.3 |
| CaCO<sub>3</sub> | 0.5 |
| CaHPO<sub>4</sub> | 0.5 |
| Salt | 0.5 |
| Vitamin-mineral premixa | 0.2 |
| Chemical composition, % of DM |  |
| DM | 47.1 |
| NDF | 36.1 |
| ADF | 20.3 |
| CP | 18.5 |
| EE | 4.1 |
| Ash | 6.6 |

<sup>a</sup>Vitamin-mineral premix (per kg): 600 mg of Mn, 950 mg of Zn, 450 mg of Fe, 650 mg of Cu, 35 mg of Se, 45 mg of I, 15 mg of Co, 450 mg of nicotinic acid, 800 mg of vitamin E, 50 kIU of vitamin D, and 115 kIU of vitamin A.

## Notes

### Competing Interest Statement

The authors have declared no competing interest.

